# Genomic legacies of ancient adaptation illuminate the GC-content evolution in bacterial genomes

**DOI:** 10.1101/2022.04.02.486805

**Authors:** Wenkai Teng, Bin Liao, Mengyun Chen, Wensheng Shu

**Affiliations:** State Key Laboratory of Biocontrol, Guangdong Key Laboratory of Plant Resources and Conservation of Guangdong Higher Education Institutes, School of Life Sciences, Sun Yat-sen University, Guangzhou 510275, China; School of Life Sciences, South China Normal University, Guangzhou 510631, China

## Abstract

In conventional views, the bacterial adaptation is characterized by strong purifying selection as well as rapid evolution in changing environments. However, the genomic GC content varies greatly but has some degree of phylogenetic stability. Using 11,083 representative genomes, we report a phylogenetically constrained bimodal distribution of the genomic GC. Results suggest that such divergence of the genomic GC can be well explained by the DNA replication and repair (DRR) system, in which multiple pathways are observed correlated to the genomic GC. The biased conservations of various stress-related genes especially the DRR-related ones imply distinct adaptive evolution of the ancestral lineages of high or low GC clades which may be induced by major environmental changes in early evolution. Furthermore, our findings support that the mutational biases resulted from these legacies of adaptation have changed the course of adaptive evolution in bacteria thus causing great variation in the genomic GC. This study demonstrates the importance of indirect effects from natural selection which may be easily misinterpreted as neutral processes.

## INTRODUCTION

Bacteria are usually supposed to be subject to strong level of purifying selection in light of huge effective population sizes (*1*). The situation hence can become confusing when understanding how bacteria have generated great variation in their genomic base composition (i.e., GC content) during the adaptive evolution (*2*). The genomic GC content of bacteria, which can vary enormously from below 20% to nearly 75%, is not normally distributed at odds with the expectation from an entirely stochastic model (*3, 4*). Therefore, the variation in genomic GC has been related to the selection acting directly on the DNA or protein sequences (*5, 6*). When bacteria are exposed to environmental stresses such as the oxidative stress, heat stress and nutrient limitation (e.g., nitrogen or carbon limitation), chemical property differences of the base pairs or amino acids likely influence the genomic GC, provided that the fitness differences are large enough (*7–9*). The correlations detected between the genomic GC and a few selective agents to some extent, though often insufficiently, favor an environmental constraint on the base composition (*10–14*).

In conventional views, bacteria evolve rapidly in response to environmental changes leaving most genetic variations non-persistent (*15*). However, previous studies have revealed remarkable phylogenetic inertia in the genomic GC, hinting at a certain level of stability of the base composition despite ever-changing environments over long timescales (*13, 16*). Thus, some attention has shifted from direct selection towards the effect of genetically conserved components, the DNA replication and repair (DRR) proteins, which potentially cause mutational biases and introduce intricate effects on the genomic GC (*17, 18*).

Bacteria suffer various DNA damages at any moment during growth especially under environmental stresses (*19*). For example, the cytosine in DNA can be deaminated spontaneously into uracil and the guanine is susceptible to oxidative damage (*20, 21*). These lesions could give rise to different types of mutations in DNA replication (DR) and directly influence the base composition (*22*). Nevertheless, multiple pathways can be recruited to cope with the DNA damages, including the base excision repair (BER), the nucleotide excision repair (NER), the non-homologous end joining (NHEJ), the mismatch repair (MMR), the homologous recombination (HR) and the translesion synthesis (TLS) (*23*). A particularly striking case is the BER in which diverse lesions including the mutagenic uracil and oxidized guanine in DNA are recognized and removed by DNA glycosylases (*22, 24*). These proteins involved in DNA repair are critical for response to environmental stresses and deletion of them could contributes to an elevated rate of related mutations (*25*). To date, several DRR-related proteins have been considered to be responsible for the variation in genomic GC (*18, 26–28*). For instance, the error-prone polymerases involved in TLS can trigger various replication errors in the newly synthesized strand affecting the mutagenic spectrum (*29–31*). However, it remains enigmatic how the DRR system has evolved in the bacterial adaptation to drive the genomic GC which correlates with the environment as well (*13*).

As an essential feature, the GC content can be related to many aspects of the genome especially the amino-acid and codon usage due to high coding density and, therefore, a better understanding for it will profoundly promote the comprehension of bacterial evolution (*32, 33*). In order to clarify the evolutionary mechanisms of the genomic GC, here we collected all currently available bacterial representative genomes. The distribution pattern and phylogenetic history of the genomic GC as well as correlated genome functions were investigated. Our results suggest an indirect way of the natural selection in which the ancient adaptations have transformed the bacterial genome (especially the DRR system) contributing to a bimodal distribution pattern of the genomic GC.

## RESULTS

### Early evolution contributes to a bimodal distribution of the genomic GC

Among 11,083 high-quality bacterial representative genomes (data S1, fig. S1), the genomic GC content varies greatly from about 16% to 77% and surprisingly displays a bimodal distribution pattern (Fig. 1A). This indicates a remarkable divergence of the genomic GC as most genomes have GC content below 50% or above 55% (Fig. 1A). Scanning the genomic GC across different taxonomic levels shows a strong level of phylogenetic inertia and particularly, more than 60% of the total variance is explained at the phylum level (Fig. 1B). The range of within-taxon variation of the genomic GC decreases gradually from the phylum to the genus level (fig. S2). Importantly, such bimodal pattern with 50% in the middle nearly disappears at the phylum level (fig. S2). Distributions of the genomic GC in most phyla are approximatively unimodal with their peak values greater or less than 50% (defined as high or low GC clade, respectively), especially in those having plenty of representatives, i.e., Proteobacteria, Firmicutes, Actinobacteria and Bacteroidetes (Fig. 1C, D, fig. S2). Results above suggest that early evolution of bacteria contributes much to the divergence of the genomic GC, which leads to the bimodal distribution.

**Fig. 1.**
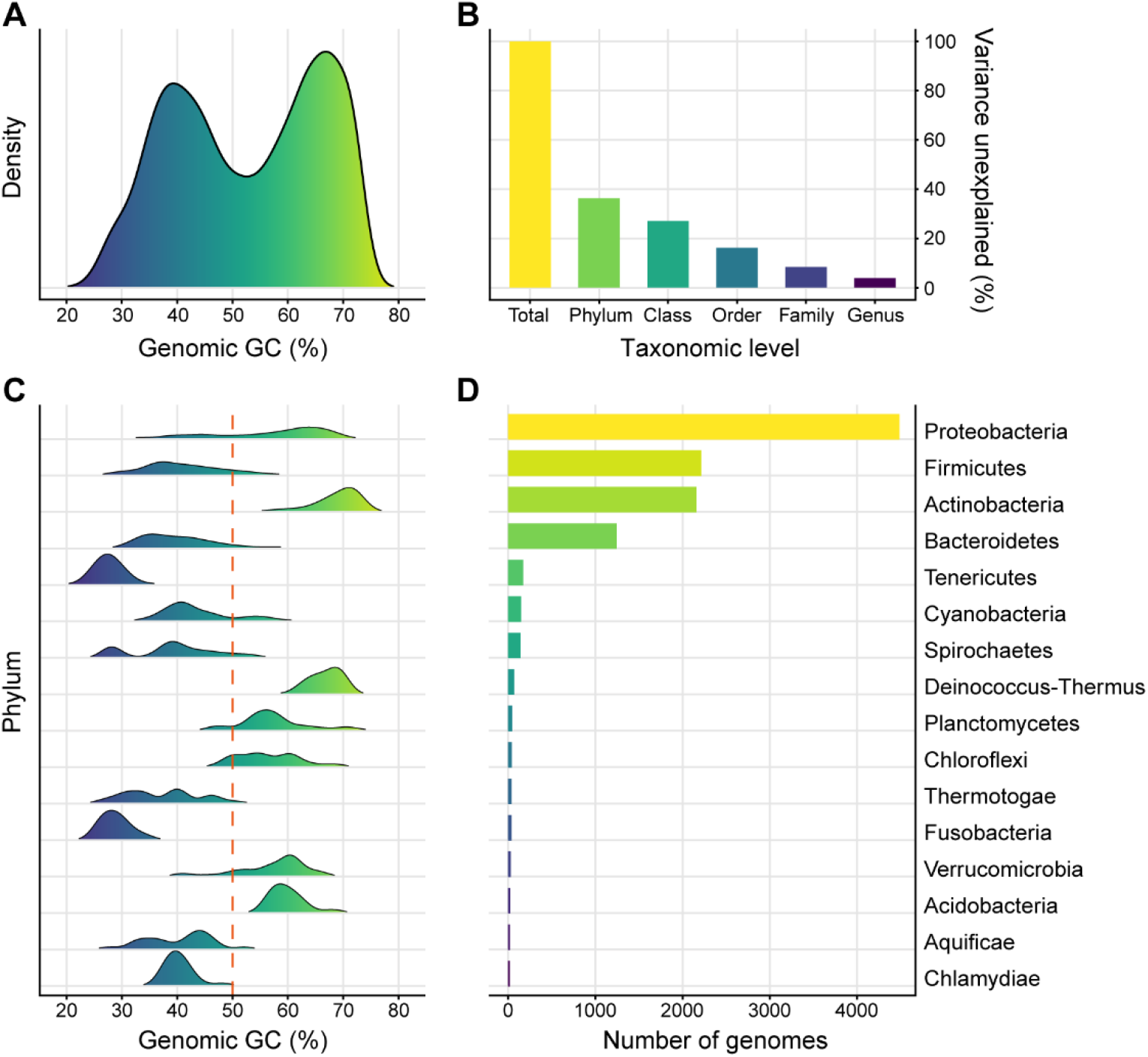
Distribution pattern and phylogenetic inertia of the genomic GC-content. (**A**) Distribution of the genomic GC-content of 11,083 bacterial representative genomes. (**B**) Variance of the genomic GC unexplained by bacterial taxonomy at different levels. (**C**) Distribution of the genomic GC of the phyla with more than 20 representative genomes. (**D**) Number of representative genomes of phyla shown in (**C**).

Ancestral state reconstruction further reveals multiple shifts in the genomic GC across the evolution of bacteria. Low GC clades are actually interspersed among high GC clades on the phylogenetic tree even within the same bacterial group (Fig. 2). In detail, while Firmicutes, Tenericutes and Cyanobacteria have relatively low genomic GC in the Terrabacteria group, clades including Actinobacteria, Deinococcus-Thermus and Chloroflexi possess genomes with comparatively high GC content (Fig. 1C, Fig. 2). When it comes to Proteobacteria, the genomic GC in most lineages especially those from Alphaproteobacteria, Betaproteobacteria and Deltaproteobacteria is rather high (Fig. 2). Despite that, low GC clades are embedded inside this group as well, taking the Epsilonproteobacteria as an example (Fig. 2, fig. S3). Particularly, the genomic GC has diverged multiple times in the Gammaproteobacteria clade which has a huge phylogenetic diversity and displays an analogous bimodal distribution (Fig. 2, fig. S3). Furthermore, the situation in the clade containing the FCB and the PVC group is also similar, where Bacteroidetes and Chlamydiae have distinct genomic GC from their relatives Planctomycetes and Verrucomicrobia (Fig. 1C, Fig. 2). Collectively, these results suggest a likely parallelism of the GC-content divergence of bacterial genomes during the early evolution of phyla or classes and also in the subsequent evolution of certain clades.

**Fig. 2.**
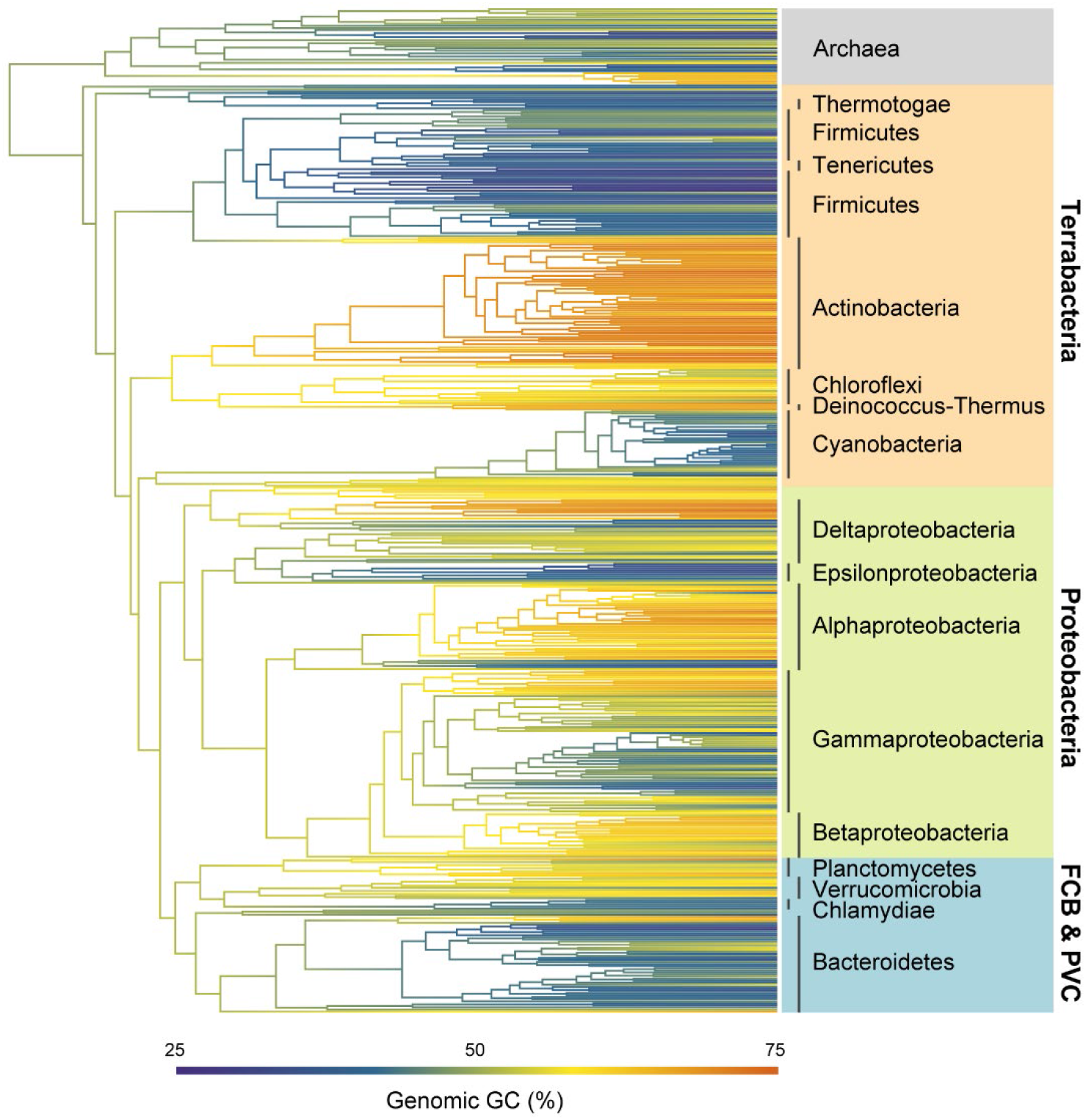
Ancestral state reconstruction of the genomic GC-content. A phylogenetic tree of the representatives of 450 bacterial families with archaeal genomes as the outgroup are used in the analysis. Divergence times were estimated using the R package APE. The genomic GC-content of ancestral nodes are reconstructed based on the average genomic GC of each family.

### DNA replication and repair explain the major variation in genomic GC

Some of previous studies suggested direct selection upon the usages of amino acids and synonymous codons which potentially influence the genomic GC (*8, 34*). Indeed, strong correlations between them are recovered in this study as well (fig. S4A, B). However, there are several findings challenge this view. Firstly, the correlation between the genomic GC and the usage of an amino acid depends on the average GC of its codons other than chemical properties generally targeted by selection (fig. S4C, D). Secondly, the genomic GC, GC content of coding sequences (*GC*_CDS_), GC content of non-coding sequences (*GC*_NCS_) and GC content contributed by the usage of amino acids (*GC*_AA_) and codons (*GC*_Codon_) are all highly correlated with each other (fig. S5). Assuming a consistent environmental impact on the whole genome via direct selection can be confusing (*5*). Finally, the phylogenetic inertia of the genomic GC as shown above highlights the view of other studies that the DRR system mainly drive the variation in genomic GC and synchronously influencing the amino-acid and codon usage (*18, 28, 35*).

As anticipated, a linear model taking all DRR-related proteins into consideration can explain up to 85% of the total variance (that is, with a multiple correlation coefficient of 0.92) suggesting that the DRR system can serve as a good predictor of the genomic GC content (Fig. 3A). Good agreement between the difference in genomic GC and the allowed minimum distance in DRR further reveals not only the sufficiency but also the necessity of the DRR system (Fig. 3B). Unlike previous scenarios, we find that the DRR-related KOs are widely implicated in the explanation of the variation in genomic GC (Fig. 3C). Among these, DnaE2, an error-prone TLS polymerase, and MutS2, a homologue of MMR protein MutS, have the highest positive and negative correlation respectively (Fig. 3C). The overwhelming majority of bacterial genomes selectively keep the DnaE2 or the MutS2 roughly according to their genomic GC (Fig. 3D).

**Fig. 3.**
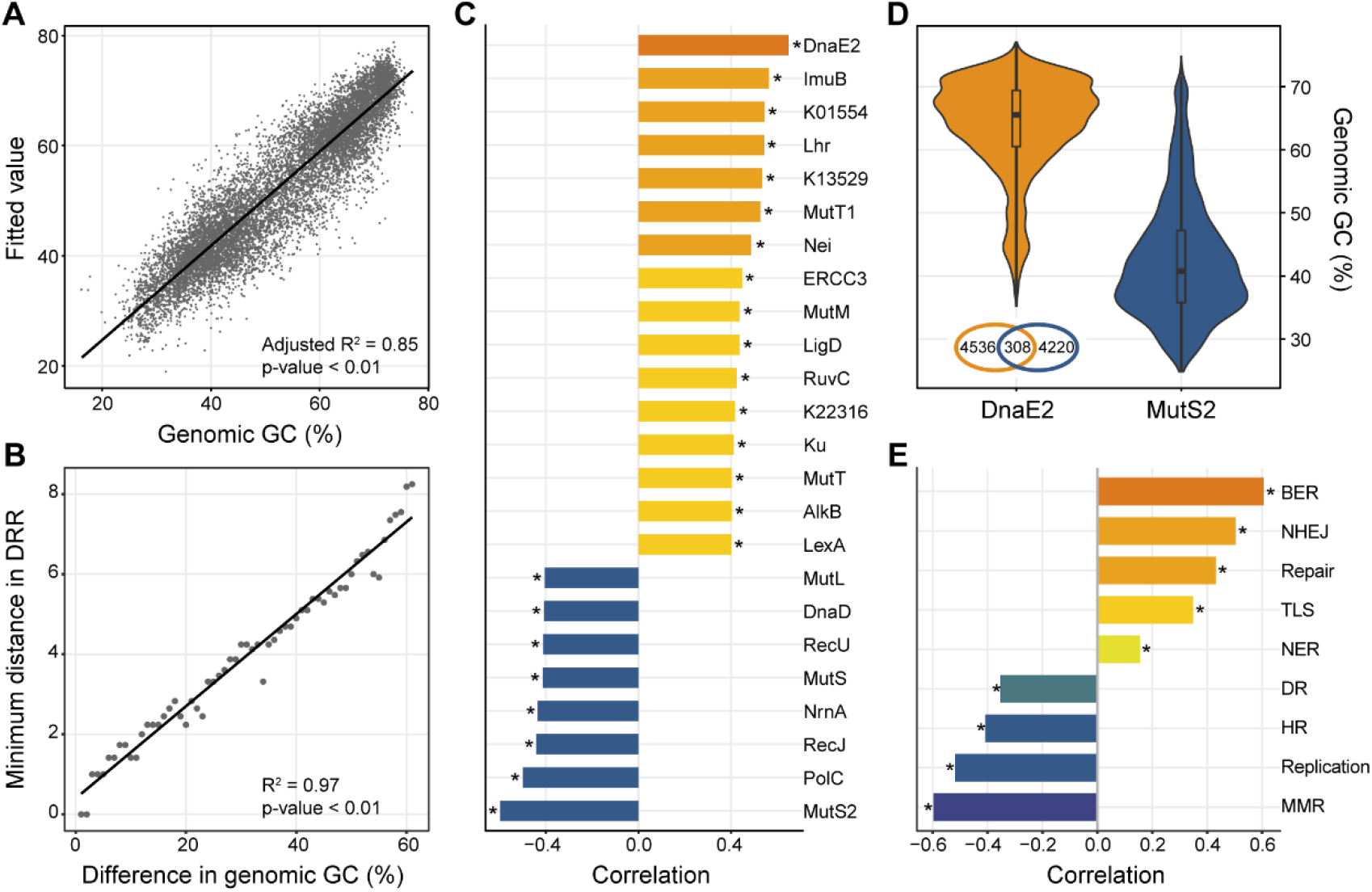
Relationships between the genomic GC-content and DNA replication and repair (DRR) system. (**A**) Multiple regression analysis of the genomic GC and DRR-related KOs. Fitted values of the genomic GC-content are shown in contrast to real values. (**B**) Minimum distance of the DRR changes with changing differences in the genomic GC. (**C**) Correlations between the genomic GC and DRR-related KOs. Only KOs with a considerable (Pearson’s r > 0.4 or r < −0.4) correlation are shown. (**D**) Comparison of the GC content of genomes containing DnaE2 and (or) MutS2. The Venn diagram inside the graph shows the number of genomes containing DnaE2 and (or) MutS2. (**E**) Correlations between the genomic GC and DRR-related pathways (DR: DNA replication; BER: base excision repair; NER: nucleotide excision repair; MMR: mismatch repair; HR: homologous recombination; NHEJ: non-homologous end-joining; TLS: translesion synthesis; Replication: DNA replication proteins; Repair: DNA repair and recombination proteins). Asterisks in (**C**) and (**E**) indicate adjusted p-value < 0.01.

Pathway analysis indicates that the positively correlated proteins are mainly involved in BER (MutM, Nei and K13529), NHEJ (Ku and LigD), TLS (DnaE2 and ImuB), NER (ERCC3) and sanitization of the nucleotide pool (MutT, MutT1 and K01554) (Fig. 3C). Many of these proteins have been shown to be crucial for growth under environmental stresses. For instance, both the MutM and Ku play important roles in combating heat as well as oxidative stresses (*25, 27*). Beyond that, the MutT and its analogue MutT1 participate in the detoxification of oxidized guanine nucleotides (*21, 36*). TLS polymerases are also important for growth under harsh environments such as starvation and high temperatures (*29*). In stark contrast, some of the MMR- and HR-related proteins including MutS, MutL, RecJ, RecU and aforementioned MutS2 are negatively correlated with the genomic GC (Fig. 3C). Overall, the DRR system displays a distinct variation pattern of positively correlated BER, TLS, NHEJ and NER versus negatively correlated DR, HR and MMR (Fig. 3E). And, in agreement with the inferred parallel divergences of the genomic GC, such pattern can also be observed individually in the Terrabacteria, Proteobacteria and FCB, PVC group (Fig. 2, fig. S6).

### Other genomic legacies correlated with the divergence of the genomic GC

In order to better understand the evolutionary scenarios, a genome-wide investigation in functional variation was conducted. Results demonstrate that plentiful proteins with various functions correlate with the genomic GC with coefficients even larger than the DRR-related DnaE2 (Fig. 4A). Similar to the situation in DRR, the highly correlated proteins can be related to the bacterial response to environmental changes of multiple factors comprising the oxygen, temperature, nutrition and osmotic pressure (Fig. 4A, table S1). For instance, YbbN, a molecular chaperone of the β-clamp, can promote the bacterial acclimation to high temperatures and are mainly conserved in the high-GC bacteria (fig. S7). Another positively correlated protein named OtsA involves in the synthesis of trehalose in response to heat, cold and osmotic stresses (Fig. 4A, table S1). Besides that, the FNR, SURF1 and YgfZ which play important roles in aerobic growth also positively correlate with the genomic GC. By contrast, several proteins including PepT, ComEB and PfkA which are required for the utilization of diverse nutrient sources and anaerobic regulatory proteins FLP and Rny negatively correlate with the genomic GC. These correlations reveal a much close relationship between the evolution of the genomic GC and that of the genome function (Fig. 4B).

**Fig. 4.**
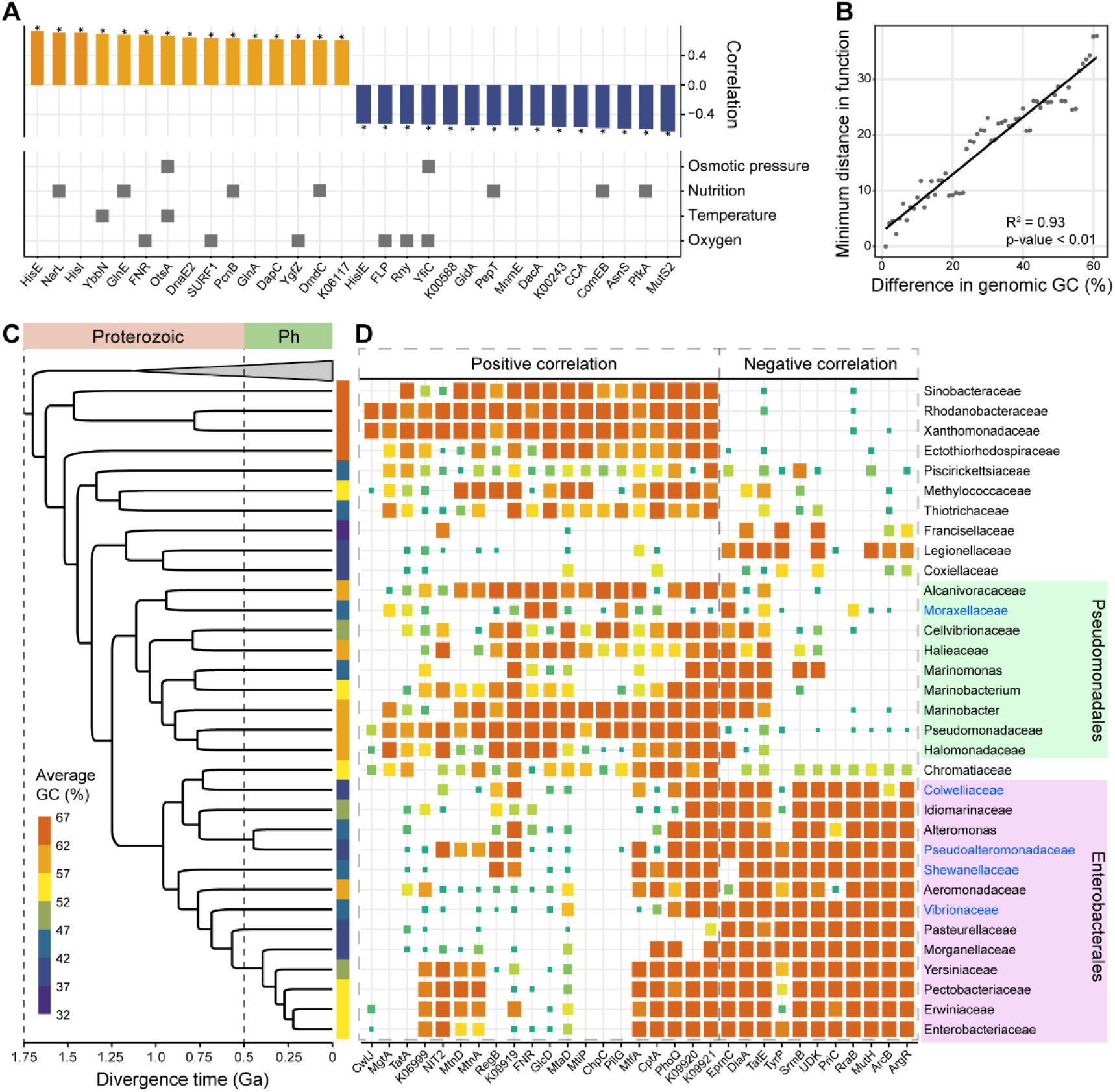
Relationships between the genomic GC-content and genome function. (**A**) Highly correlated KOs and their involvement in bacterial response to the changes in four environmental factors. Only KOs with a considerable (Pearson’s r > 0.6 or r < −0.5) correlation are shown. Asterisks indicate adjusted p-value < 0.01. (**B**) Minimum distance of the genome function changes with changing differences in the genomic GC. (**C**) Phylogenetic tree of the lineages in Gammaproteobacteria with estimated divergence times (Ga: billion years ago). Genomes belonging to Betaproteobacteria are used as the outgroup. Color strips beside the tree reflect the average genomic GC-content. The names of lineages with presence of more than 5 cold-adapted species are shown in blue. (**D**) Conservation of the protein within each major lineage belonging to the Gammaproteobacteria. Only KOs with a considerable (Pearson’s r > 0.5 or r < −0.45) correlation are shown. Positively correlated KOs and negatively correlated KOs are indicated using dashed boxes.

Further analysis indicates that parallel gene loss likely contributes substantially to the biased conservations of functional proteins. First, most highly correlated proteins can be observed strongly conserved in distantly related lineages (fig. S8-9). For example, positively correlated KOs including DnaE2, ImuB, MutM, MutT, FNR and YbbN are highly conserved in Actinobacteria, which belong to Terrabacteria, as well as Alpha- and Beta-proteobacteria (fig. S8-9). Thus, the related genes should have been present in the last common ancestor (LCA) of the two groups and lost in parallel during later evolution. Further, some of the highly correlated KOs such as MutT and Lhr are even pretty conserved in archaea suggesting much earlier origins of their genes (fig. S10). Second, there are much more proteins positively correlate with the genomic GC, which implies a severer loss of functional genes in low GC clades (fig. S11). Consistent with this, the bacterial genome size in general positively correlates with the genomic GC (fig. S12). Such loss of the positively correlated functional genes is particularly striking for the low GC lineages in Gammaproteobacteria, where the divergence of the genomic GC appeared much more recently (Fig. 4C, D). Notably, the highly correlated KOs are either pretty conserved or almost entirely lost in a certain lineage, indicating that they are genomic legacies from early evolution (fig. S13).

### Bacterial evolution and major environmental changes in early history

According to the molecular dating of the bacterial tree, multiple important divergence events of the genomic GC likely occurred during the period from the mid-Archaean to the Paleoproterozoic (i.e., early stages of the evolution of major clades), accompanied by the loss of many conserved genes (Fig. 2, Fig. 5). As revealed in the previous study, there was a peak genetic innovation of bacterial ancestors named the Archaean genetic expansion and rapid gene loss followed it in different lineages (*37*). This is in line with our findings (Fig. 5). Bacterial evolution through gene loss has commonly been associated with the adaptation to new environments (*38–40*). Biased conservations of the functional genes between high and low GC clades suggests adaptation to different environmental conditions mainly arising from differential heat and oxidative stress (Fig. 3C, Fig. 4A). Further, the similar evolutionary pattern of the DRR system as demonstrated above implies a similar mechanism of adaptation in high or low GC bacteria from different groups (Fig. 3E, fig. S6).

**Fig. 5.**
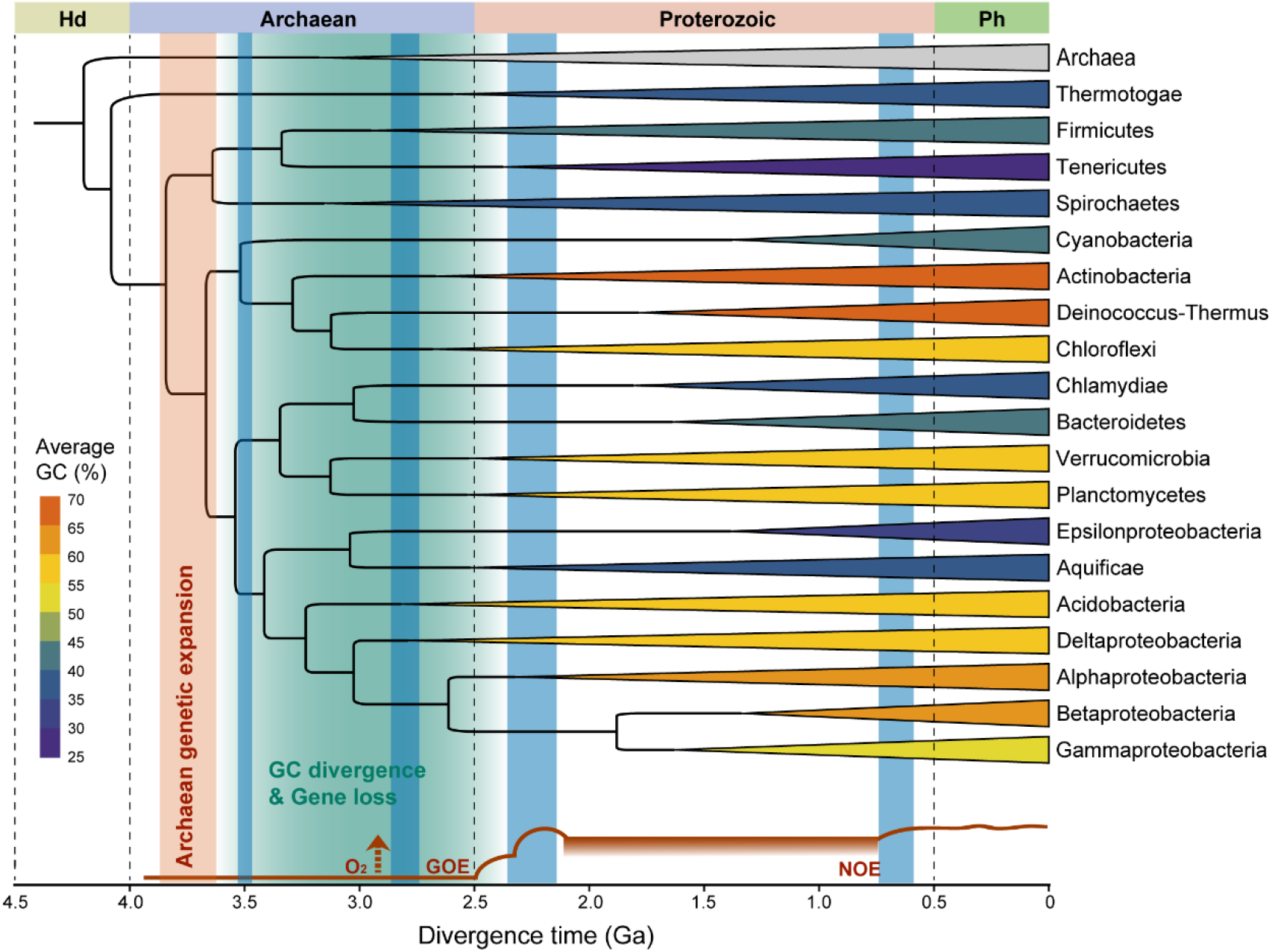
Molecular clock analysis of major bacterial clades. Divergence times were estimated using the MCMC algorithm (Ga: billion years ago). Archaeal genomes were used as the outgroup. Clades are colored according to their average genomic GC. The green shaded area indicates the time period when the divergence of the genomic GC and gene loss might have occurred. The dark red bar indicates the inferred Archaean genetic expansion (*37*). The lower curve (in dark red) indicates the evolution of the atmospheric oxygen over Earth history (GOE: Great Oxidation Event, NOE: Neoproterozoic Oxygenation Event) (*41*). Time periods for glaciation before the Phanerozoic are denoted by blue bars (*45*).

The Earth’s history is truly characterized by dramatic changes in atmospheric oxygen and temperature (*41–43*), represented by two oxidation events and a series of glaciation events (Fig. 5). Although the timing of the first emergence of oxygen remains highly uncertain (Fig. 5), molecular analysis has revealed that all major bacterial clades except for the Thermotogae descend from a universal oxygen ancestor before the Great Oxidation Event (*44*). Based on the loss of various stress-related genes and the conservations of PepT, ComEB and PfkA, the ancestral lineages of low GC bacteria likely have experienced strong selection by oligotrophic environments with low temperature and oxygen. This is consistent with the current knowledge about Archaean climate (Fig. 5) that though mostly temperate to hot, the earliest evidence for the glaciation can date back to early Archaean (*45*).

To further investigate this relationship, we collected cold-adapted species, including the defined psychrophilic and psychrotolerant bacteria, which have been isolated from the similar environments such as ocean depths, polar regions and subglacial sediments (data S2). As anticipated, both the psychrophilic and psychrotolerant bacteria show skewed bimodal distribution of the genomic GC, hinting at a possible association between the history of cold adaptation and genomic GC reduction (Fig. 6A, fig. S14). These species mainly belong to Proteobacteria (especially Gammaproteobacteria) and Bacteroidetes and particularly, 25% of them exactly come from low GC lineages in Gammaproteobacteria (Fig. 4C, Fig. 6B). Such reduction of the genomic GC can also be detected within each clade (Fig. 6C). Despite the lack of information on ancient bacterial adaptation, our results still provide a glimpse into the evolution of the genomic GC in the context of major environmental changes.

**Fig. 6.**
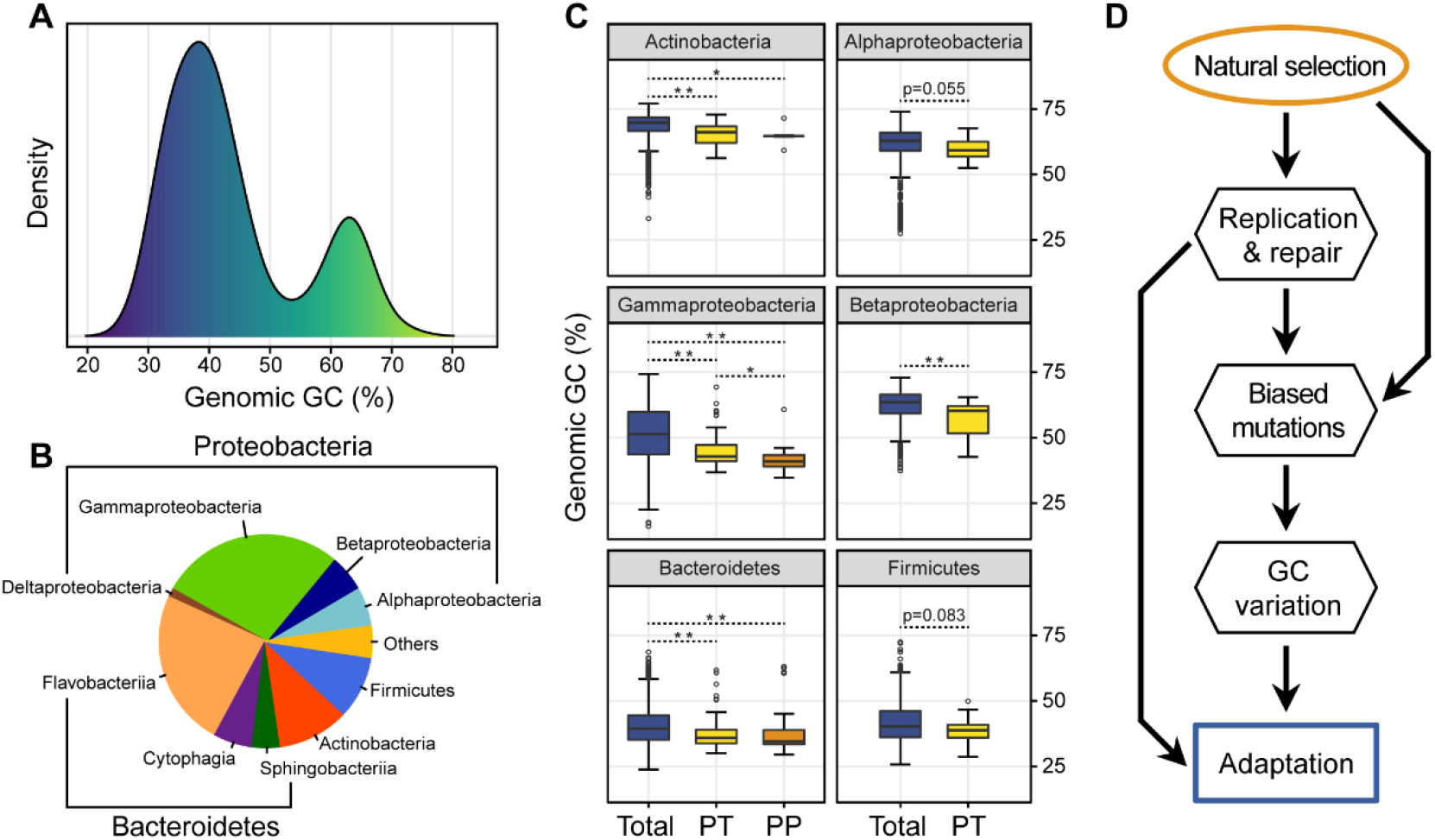
Relationship between the genomic GC and ancient bacterial adaptation. (**A**) Taxonomy composition of identified cold-adapted species. (**B)** Distribution of the genomic GC of psychrophilic species. (**C**) Comparison of the genomic GC of total, psychrotolerant (PT) and psychrophilic (PP) species within each clade. Single asterisk indicates adjusted p-value < 0.05 and double asterisk indicates adjusted p-value < 0.01. (**D**) Evolutionary mechanism of the genomic GC in bacteria.

## SUMMARY

A long-standing controversy exists regarding what mechanism accounts for the great variation in the GC content of bacterial genomes (*5, 8, 46*). Previous studies have mostly focused on the influences from environments and mutational biases caused by DRR-related proteins (*10, 12, 13, 18, 27, 28*). By performing comparative genomic analyses, we have provided more information about the evolution of the genomic GC, which potentially contributes to an improved understanding of the mechanism. Firstly, our results demonstrated a phylogenetically constrained bimodal distribution of the genomic GC. This should negate rapid evolution of the genomic base composition in changing environments. We further showed that the evolution of the DRR system mainly accounts for the variation in the genomic GC which highlights the importance of mutational biases. The observation of AT-biased mutational patterns even in some of the high GC bacteria (*17, 47*) are not contradictory to this because our findings suggest that the extant values of the genomic GC of most bacteria are principally determined by their phylogenetic history in early time windows. However, this does not mean that our results support the neutral theory or neutral fixation of the biased mutations. On the contrary, based on the fact that the genomic GC deeply influences the usage of amino acids even in highly conserved regions of the ribosomal proteins which are not likely neutral (fig. S15), natural selection is still at work during the fixation of biased mutations. All these findings lead to the viewpoint put forward in a recent study that the mutational biases can also shape the course of adaptive evolution in bacteria (*48*).

Our global comparison of the functional inventory reveals that the ancestral lineages of low-GC bacteria have undergone parallel loss of the stress-related genes especially those involved in DNA repair, implying the effects from adaptive evolution. Previous studies have reported many cases of GC-content reduction in company with genome streamlining in prokaryotes (*49–53*). Two hypotheses, genetic drift under relaxed selection and natural selection due to nutrient restriction have been developed by researchers regarding the symbiotic and free-living bacteria, respectively (*53–55*). Nevertheless, given the importance of DRR-related genes (*22, 25*), a relaxation in one direction of the environmental constraints on them should be necessary as well for their loss in free-living bacteria (*56, 57*). From a thermodynamic perspective, bacteria are less susceptible to DNA damages including the oxidative damage at lower temperatures, thus in theory requiring less repair-related genes (*58, 59*). Accordingly, reduced ability of the DNA repair and the loss of functional genes have been observed in some of the cold-adapted species (*60–64*). In Earth’s history, major glaciation events took place multiple times from early to Archaean and have deeply influenced the evolution of life (*42, 45*). It has been reported that bacteria descended from a thermophilic ancestor and adapted to lower temperatures subsequently (*65*). Therefore, the acclimation to extremely cold environments might be one of the major promoters of gene loss and GC-content divergence of bacterial genomes.

Overall, our study demonstrates that the legacies of ancient adaptation of bacteria still accounts for the major variation in the genomic GC possibly due to strong purifying selection in their subsequent evolution. Natural selection drives the genomic GC, as highlighted in a recent paper (*66*), in an indirect way where the adaptive evolution of the DRR induces biased mutations and have changed the course of bacterial evolution (Fig. 6D). This idea is plausible, for example, when considering the result that the loss of MutM and MutT, which can lead to AT-bias and GC-bias respectively under oxidative stress (*21*), can both be observed in some of the low-GC clades (Fig. S8). Naturally, the variation in genomic GC significantly influences the average amino-acid properties of proteomes, including the N/C ratio and the hydrophobicity (Fig. S16), which may in turn shape the bacterial fitness (*6, 8*). In other words, selection does not act directly on the bias of mutations or base composition, but rather on the DRR system and the resulted mutations (Fig. 6D). Our results, in accordance with many previous studies, agree with that the GC contents of microbiomes are under constraints from both phylogeny and environments (*10, 13, 27*). The molecular mechanisms underlying the effects from the DRR system on the genomic GC and especially their evolutionary history, of course, require further investigation. Still, our findings should provide more insights into the evolutionary mechanisms of the variation in genomic GC in bacteria and lay the groundwork for future studies.

## METHODS

### Genomic data collection and phylogenetic analysis

A total of 11,502 NCBI-defined reference and representative genomes, together with corresponding genes and proteins, were downloaded from the RefSeq database (https://www.ncbi.nlm.nih.gov/refseq/) on 30 March 2020. Of these, 11,083 and 419 genomes belong to bacteria and archaea respectively. Quality measures including the completeness and the contamination of genomes were estimated using CheckM (*67*). For phylogenetic analysis one genome with the highest quality was sampled from each family according to the classification in NCBI taxonomy database (https://www.ncbi.nlm.nih.gov/taxonomy/). A set of 381 markers ware extracted from the proteins of each genome and aligned using MAFFT v7.427 as previously described (*68*). All alignments were individually trimmed to remove poorly aligned positions using trimAl v1.4.15 (*69*). The trimmed alignments were then concatenated into a supermatrix. A maximum likelihood phylogenetic tree was finally constructed from the concatenated alignments using RAxML v8.2.12 under the PROTGAMMALG model, followed by 100 bootstrap replicates (*70*). In this tree archaeal genomes served as the outgroup. Using the same procedures, phylogenetic trees for *Gammaproteobacteria* and *Desulfovibrio* were also constructed with their relatives as outgroups. All trees were visualized using iTOL (*71*).

### Quantification of base composition bias

The genome size, genomic GC content and amino acid and codon abundance in all genomes were calculated using custom Python or Perl scripts. To evaluate the concordance on the genomic GC content, coding and non-coding sequences were separately calculated as *GC*_CDS_ and *GC*_NCS_. Furthermore, the GC content contributed by amino-acid usage was calculated by fixing the synonymous codon usage:

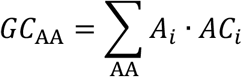

where *A_i_* is the abundance of amino acids (AA) in a genome and *AC_i_* is the average codon GC of an amino acid provided than there is no bias in synonymous codon usage. The GC content contributed by synonymous codon usage was calculated by fixing the amino-acid usage:

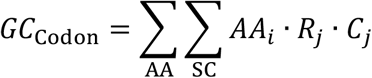

where *AA_i_* is the average abundance of amino acids in all genomes, *R_j_* is the relative abundance of a synonymous codon (SC) of an amino acid and *C_j_* is the GC content of this codon.

### Quantification of amino acid properties

To evaluate the influence of variation in amino acid composition, chemical properties including the molecular weight, VdW volume, isoelectric point (pI), hydrophobicity, polarity, number of nitrogen atoms (N number) and N/C ratio were collected via literature searches (data S3). Average properties of each bacterial proteome were calculated based on the amino acid composition (taking average hydrophobicity as an example):

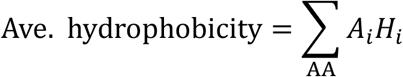

where *H_i_* is the hydrophobicity of a specific amino acid.

### Functional annotation of genomes

All genomes are annotated with KOs using the hidden Markov models database of KEGG Orthologs (KOfam, downloaded from ftp://ftp.genome.jp/pub/db/kofam/ on 14 April 2020) implemented in HMMER v3.1b2 with the following parameters: E-value ≤ 10^−5^; alignment coverage ≥ 0.7 (*72*). The KOs in KEGG pathways of DNA replication (ko03030), base excision repair (ko03410), nucleotide excision repair (ko03420), mismatch repair (ko03430), homologous recombination (ko03440), and non-homologous end-joining (ko03450) and BRITE categories of DNA replication proteins (ko03032) and DNA repair and recombination proteins (ko03400) are defined as DRR-related KOs.

### Molecular dating of phylogenetic trees

Divergence times of the above bacterial tree were firstly estimated using the R package APE, with maximum constraints of 4.52 Ga and 3.225Ga for the root and the crown Cyanobacteria, respectively (*73*). Ancestral state reconstruction of the genomic GC was carried out using the R package phytools. For a more accurate molecular clock, a reduced bacterial tree was reconstructed following the same procedures for the sake of computational expense by randomly sampling three genomes in each phylum (class in Proteobacteria) from the above bacterial tree. Three non-representative genomes belonging to the newly named Melainabacteria (assembly accession: GCA_001858525.1, GCA_002102725.1 and GCA_013216135.1) were downloaded and included in this tree (fig. S17). The divergence times were then estimated using the approximate likelihood calculation in MCMCtree v4.9 under the LG model (*74*). The node splitting *Cyanobacteria* and *Melainabacteria* was constrained to 2.5-2.6 Ga (*68*). The convergence of MCMC samples was tested in Tracer v1.7.1 (*75*). Moreover, the divergence time estimation was also performed using the *Gammaproteobacteria* tree with a maximum constraint of 2.19 Ga for the root according to the result of the bacterial tree. All time calibrated trees were visualized using FigTree v1.4.4 (https://github.com/rambaut/figtree/).

### Identification of cold-adapted species

Literature searches were conducted to collect possible cold-adapted bacterial species with the following keywords: psychrophilic, psychrotolerant, antarctic, arctic, glacier/glacial, deep-sea, cryosphere. Only species presented in our data set were considered for further analysis. We also used data from the BacDive database to obtain more information of growth temperatures (*76*). Here, psychrophilic bacteria are organisms growing well from below 5 to about 20 °C with an optima no more than 15 °C (*77*). While psychrotolerant bacteria are organisms grow optimally between 15 and 25 °C and show now growth at above 30 °C (*77*).

### Statistical analysis and visualization

Statistical analysis, including the calculation of Pearson correlation coefficient and multiple regression analysis, and visualization were mainly performed using R v4.0.3. Variance of the genomic GC unexplained by taxonomy is calculated based on the sum of squared deviations (SS):

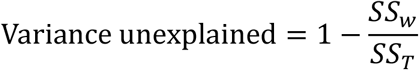

where *SS_T_* is the total SS of the genomic GC and *SS_W_* is the sum of within-taxon SS at a specific taxonomic level. Wilcoxon tests were conducted to compare intergroup differences of the genomic GC. All estimated p-values in multiple analyses were adjusted using the *Bonferroni* method. The much conserved 16 ribosomal proteins in bacteria ware extracted from the proteins of each genome as described in previous study (*78*) and then aligned and trimmed equally as described above. Finally, the variance partitioning analyses (VPA) of the amino acid composition of whole-genome proteins and ribosomal protein alignments were performed using the vegan package (https://cran.r-project.org/web/packages/vegan/index.html).

