## Supplementary material for "Genomic legacies of ancient adaptation illuminate the GC-content evolution in bacterial genomes": Figures S1 to S17, Tables S1, Data S1 to S3: Supplementary.pdf

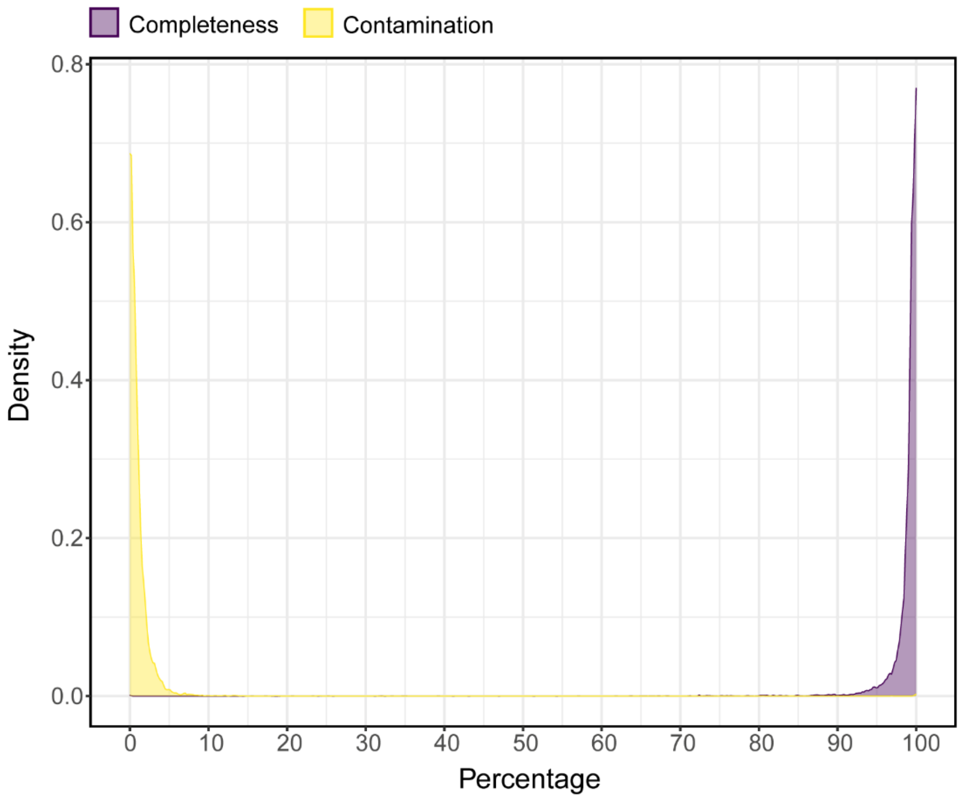


**Figure S1. Quality distribution of 11,083 bacterial representative genomes.** Purple represents genome completeness and yellow represents genome contamination.

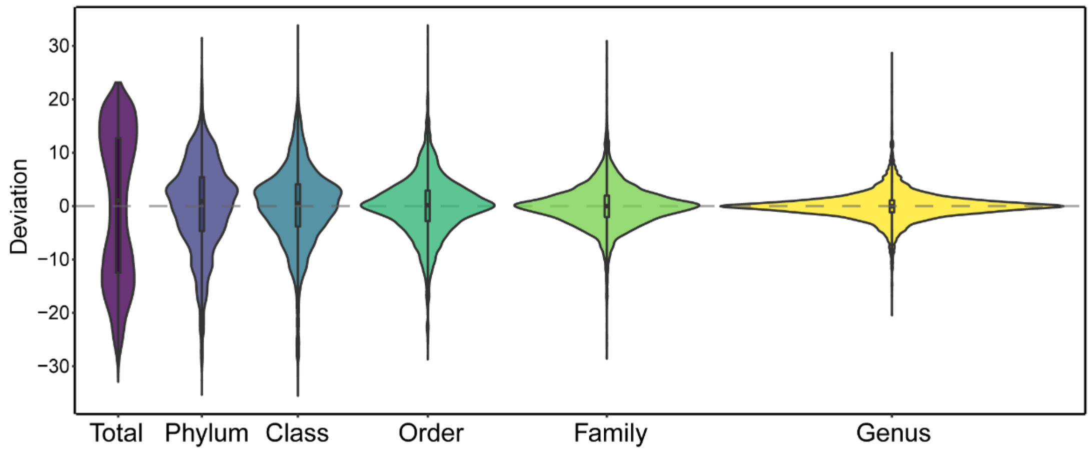


**Figure S2. Within-taxon variation of genomic GC content at different taxonomic levels.** The deviation represents the difference between GC content value and the average value within a specific taxon.

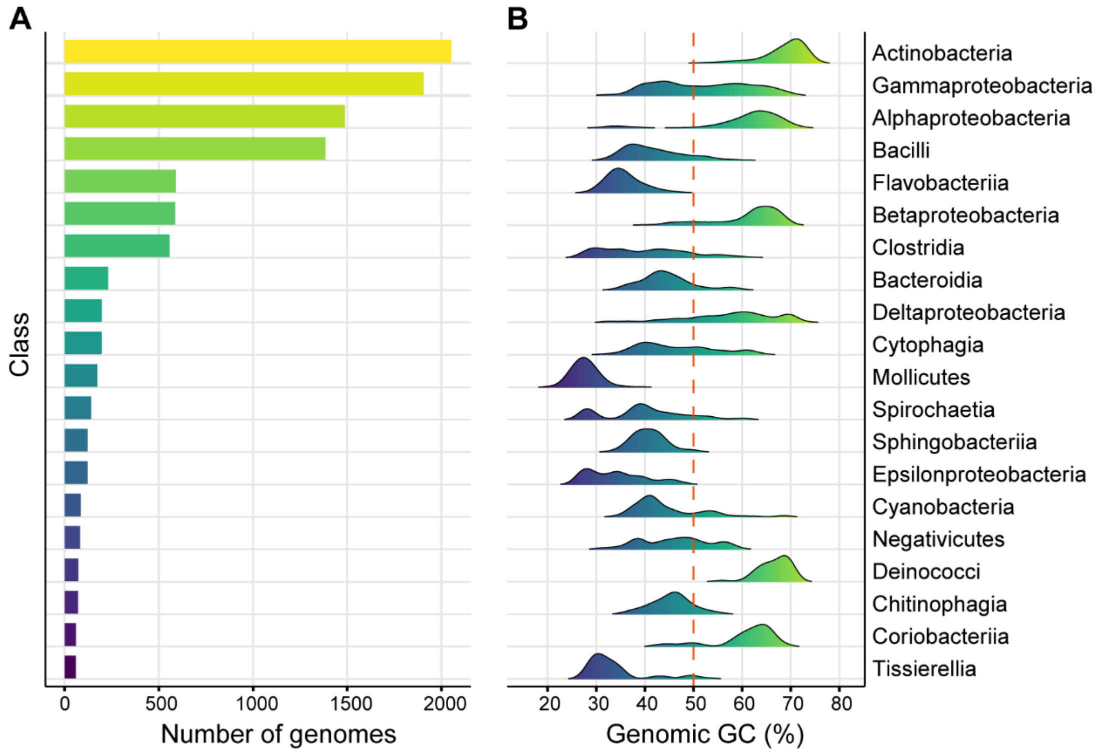


**Figure S3. Distribution of genomic GC content at the class level.** (**A**) Distribution of genomic GC of classes with more than 50 representative genomes. (**B**) Number of representative genomes of classes in (**A**).

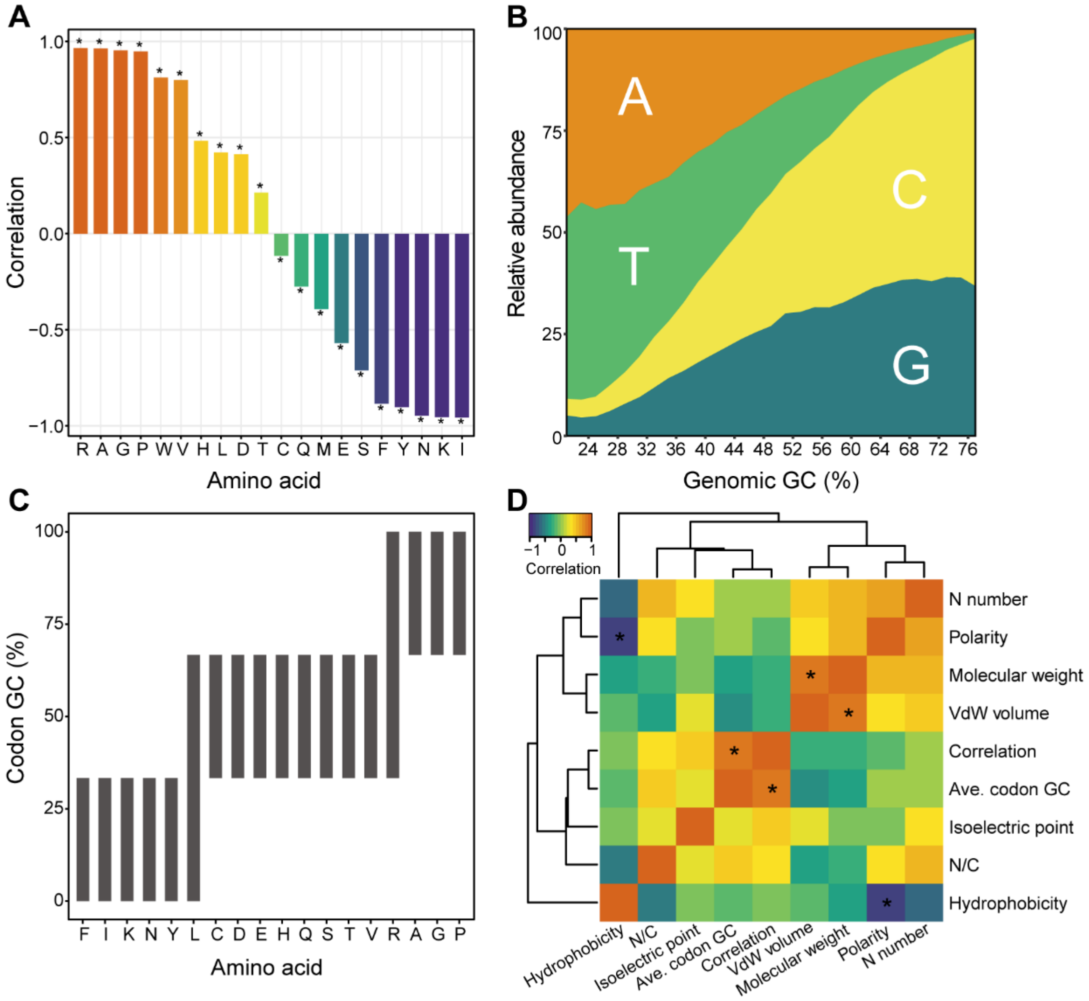


**Figure S4. Relationships between the genomic GC, codon usage and amino acid usage.** (**A**) Correlations between the genomic GC and the abundance of amino acids. (**B**) Abundance of codons with different base in the third position is correlated to the genomic GC content. Only quartets (i.e., the codons of A, G, P, V and T) are analyzed. (**C**) GC-content range of the codons of each amino acid. (**D**) Pairwise correlation tests between the correlation with the genomic GC, the average codon GC and chemical properties of each amino acid. Asterisks in (**A**) and (**D**) indicate adjusted p-value < 0.01.

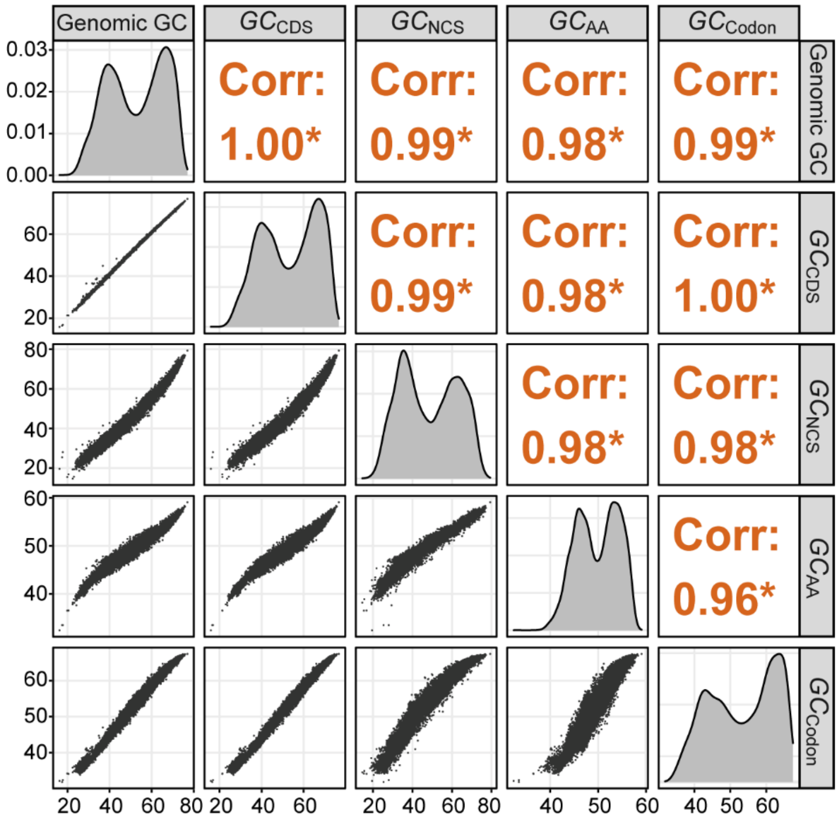


**Figure S5. High-consistency in the GC content variation.** Pairwise correlation scores between the genomic GC, GC content of coding sequences ($GC_{CDS}$), GC content of non-coding sequences ($GC_{NCS}$) and GC content contributed by amino-acid usage ($GC_{AA}$) and synonymous codon usage ($GC_{Codon}$) are shown.

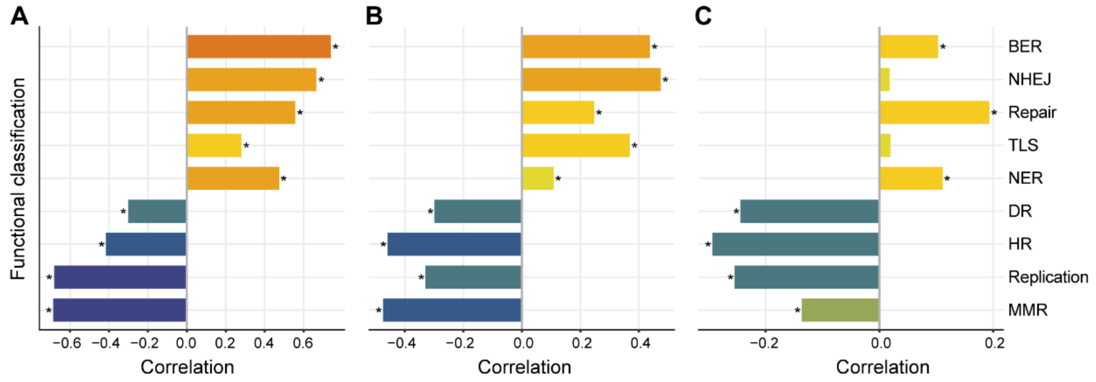


**Figure S6. Correlations between the genomic GC and DRR-related pathways.** (**A**) Correlations between the genomic GC and DRR-related pathways in the Terrabacteria clade (**B**) Correlations between the genomic GC and DRR-related pathways in the Proteobacteria clade. (**C**) Correlations between the genomic GC and DRR-related pathways in the FCB & PVC clade.

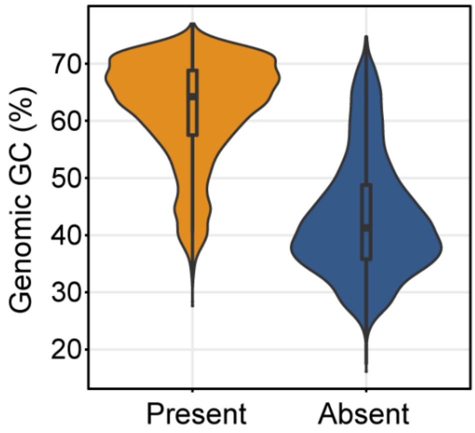


**Figure S7. Comparison of the GC content of genomes containing YbbN or not.**

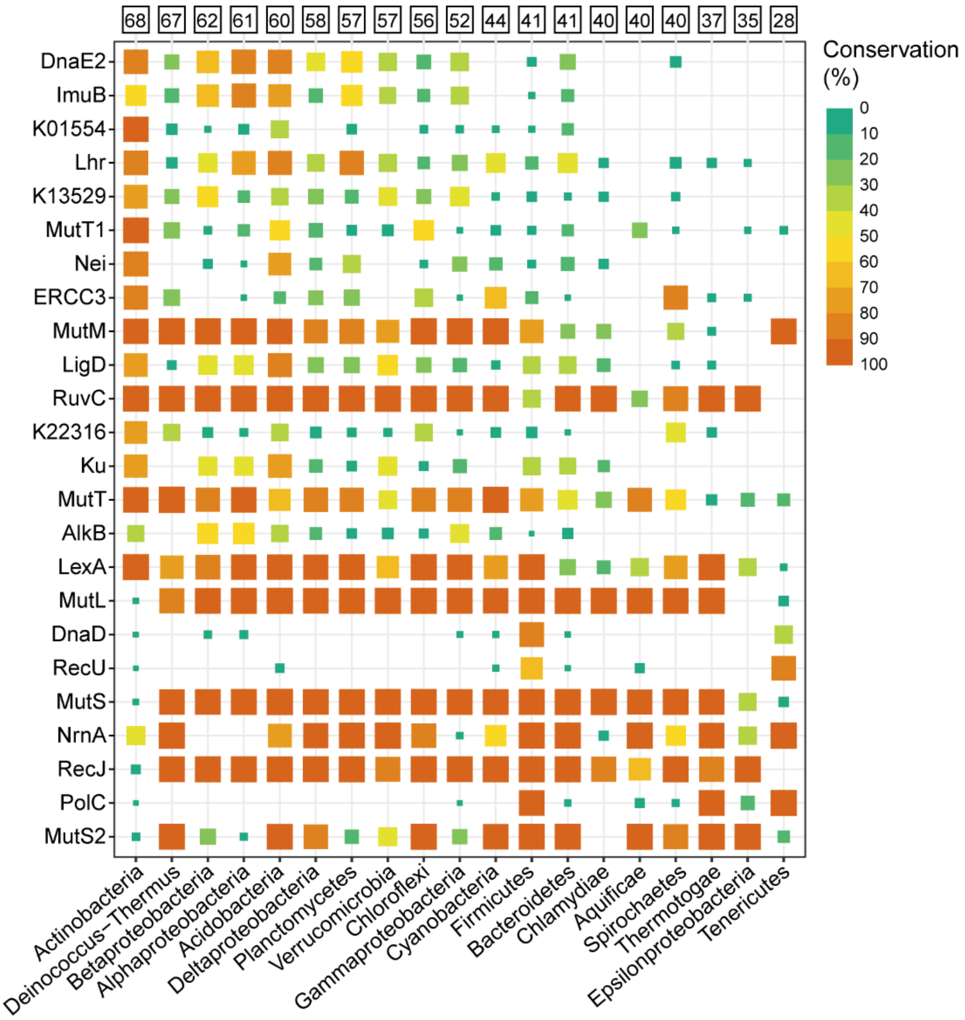


**Figure S8. Conservation of DRR-related KOs shown in Fig. 3C in major clades of bacteria.** The number in square above the plot shows the average genomic GC content of each clade.

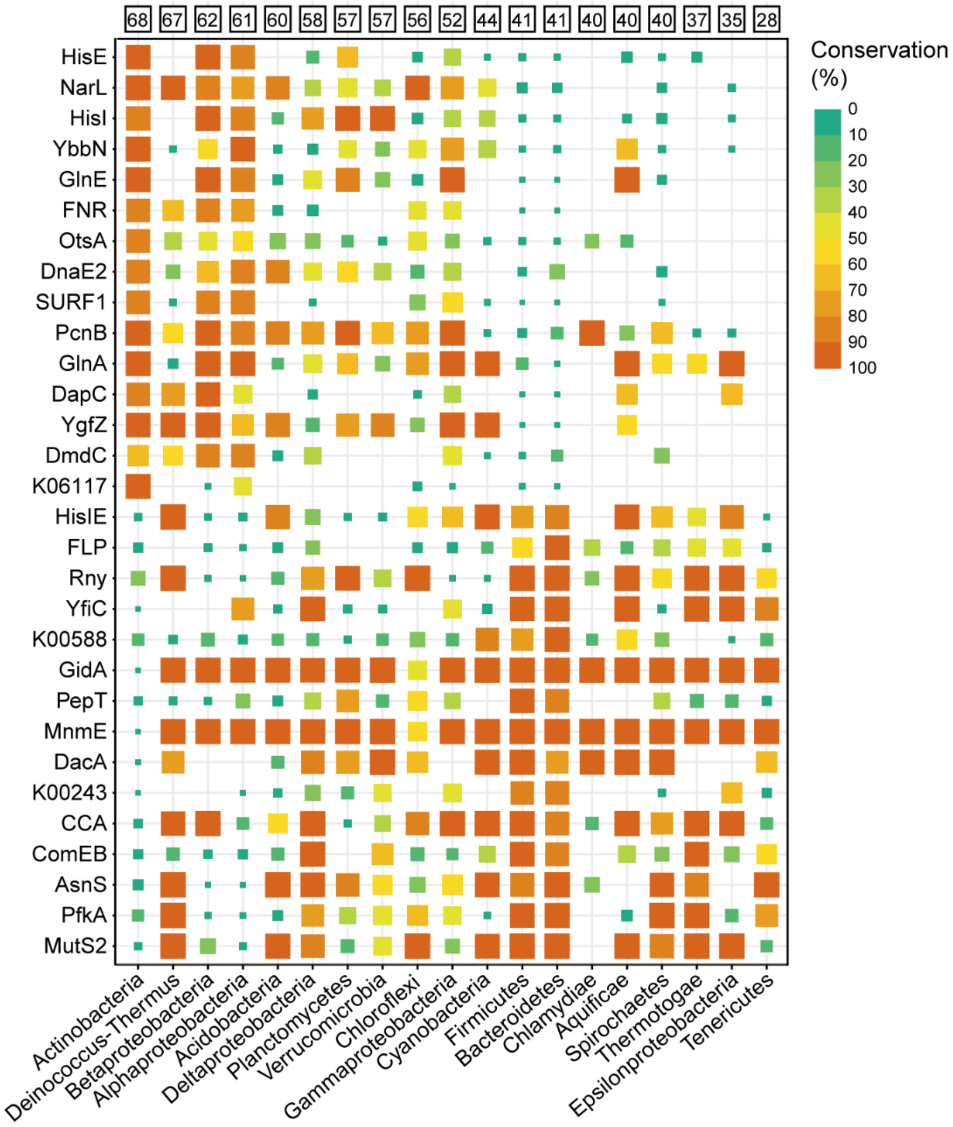


**Figure S9. Conservation of highly correlated KOs shown in Fig. 4A in major clades of bacteria.** The number in square above the plot shows the average genomic GC content of each clade.

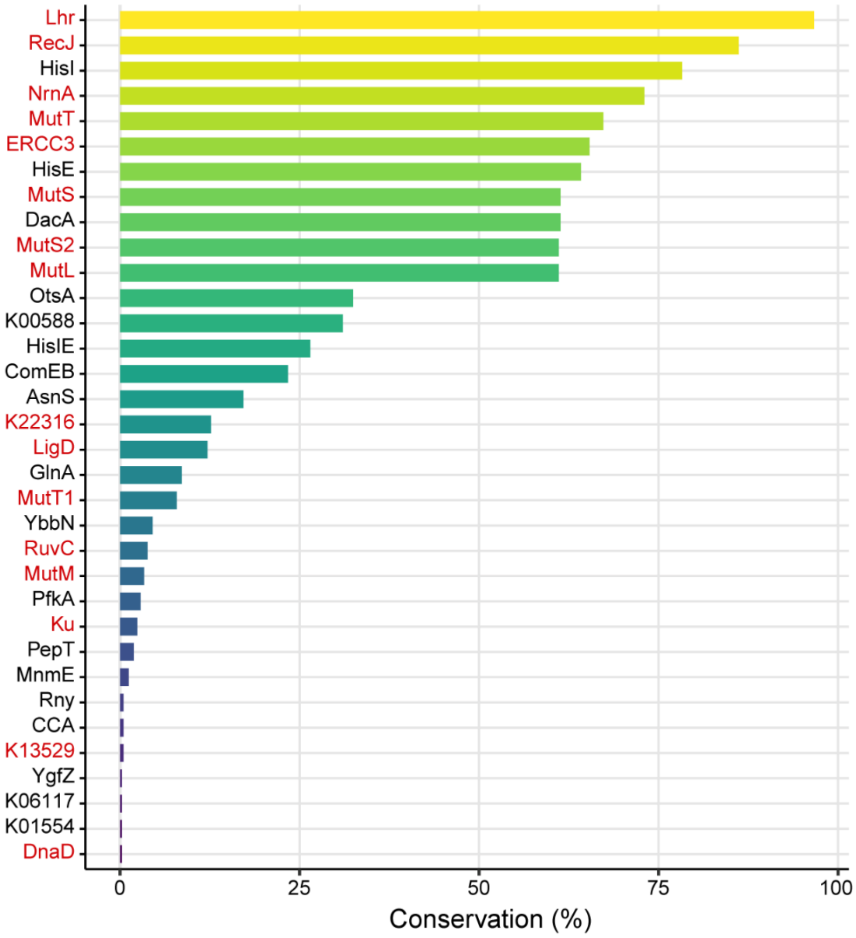


**Figure S10. Conservation of KOs shown in Fig. 3C and Fig. 4A in archaea.** The names of DRR-related KOs are shown in red. Completely missing KOs are neglected.

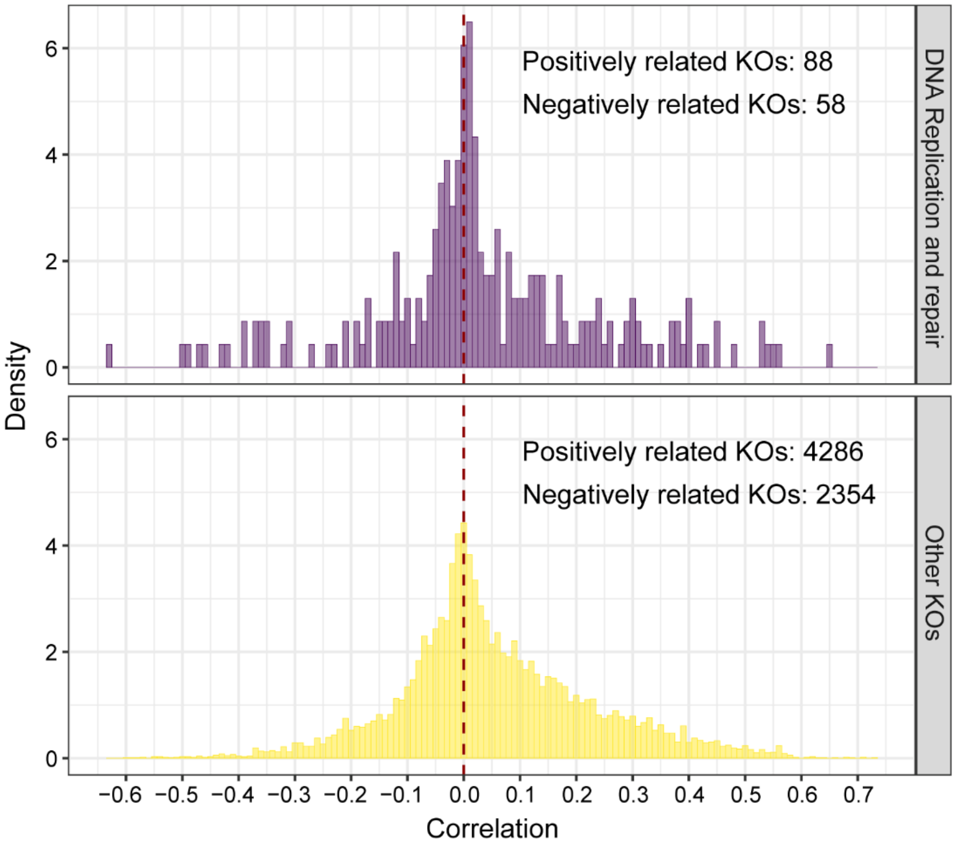


**Figure S11. Distribution of the correlations between the genomic GC content and annotated KOs.** DRR-related KOs and the others are displayed separately.

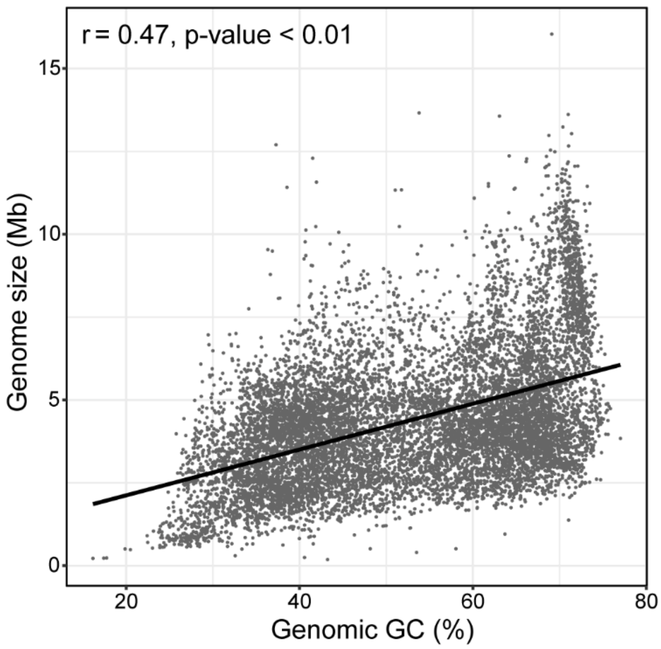


**Figure S12. Linear regression analysis between the genomic GC and genome size.**

Pearson's correlation coefficient and p-value are labeled on the plot.

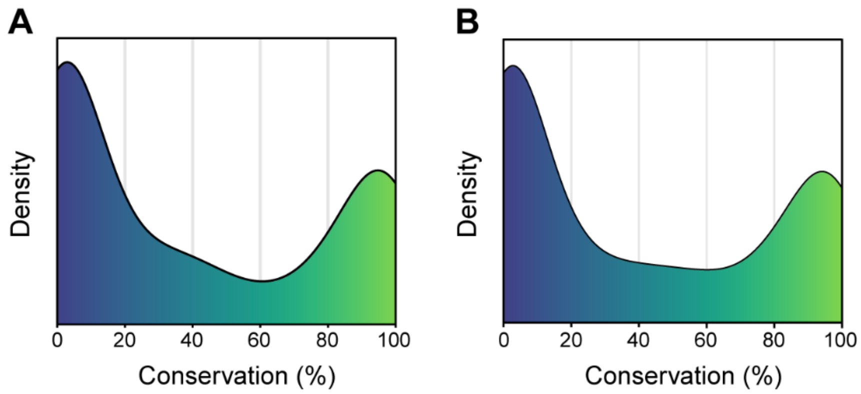


**Figure S13. Distribution of the conservation rate of highly correlated KOs.** (**A**) Distribution of the conservation data shown in Figure S8. (**B**) Distribution of the conservation data shown in Figure S9.

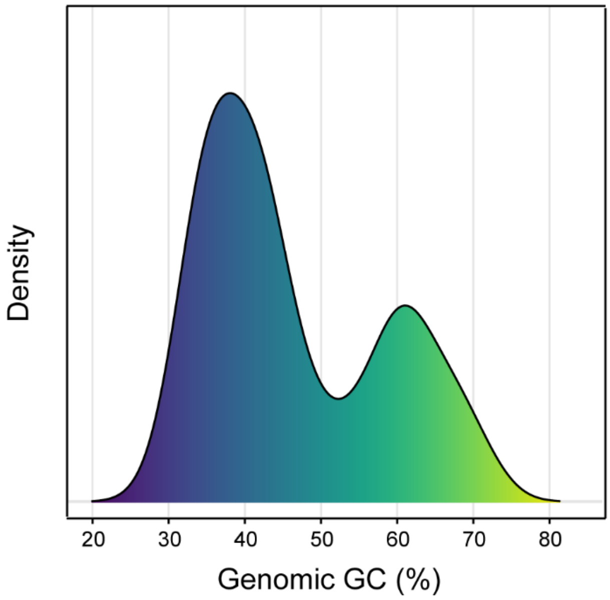


**Figure S14. Distribution of the genomic GC of psychrotolerant species.**

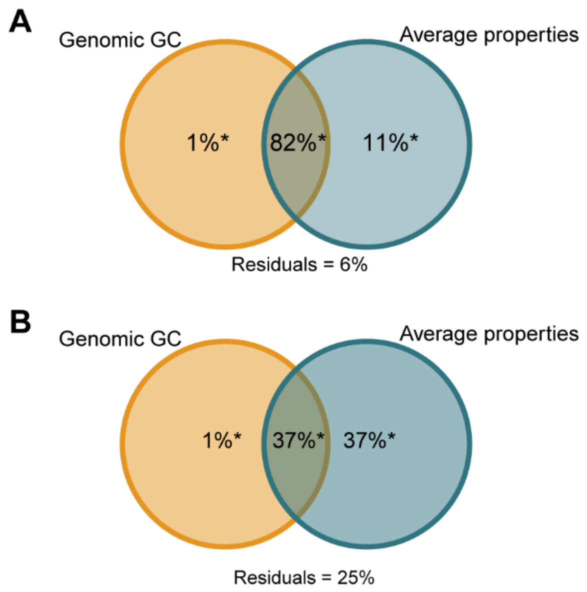


**Figure S15. Variance partitioning analysis of the amino acid composition.** (**A**) Variance partitioning analysis of the amino acid composition of all bacterial proteins. (**B**) Variance partitioning analysis of the amino acid composition of the conserved regions in 16 ribosomal proteins. Asterisks beside the percentages indicate adjusted p-value $< 0.01$.

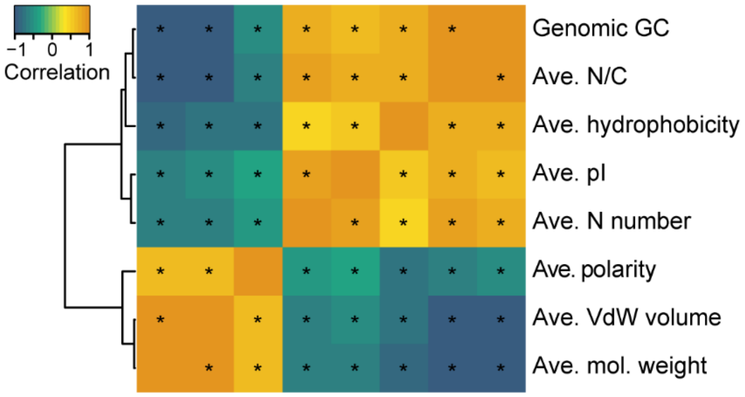


**Figure S16. Correlations between the genomic GC and average amino acid properties in bacteria.** Asterisks in the plot indicate adjusted p-value < 0.01.

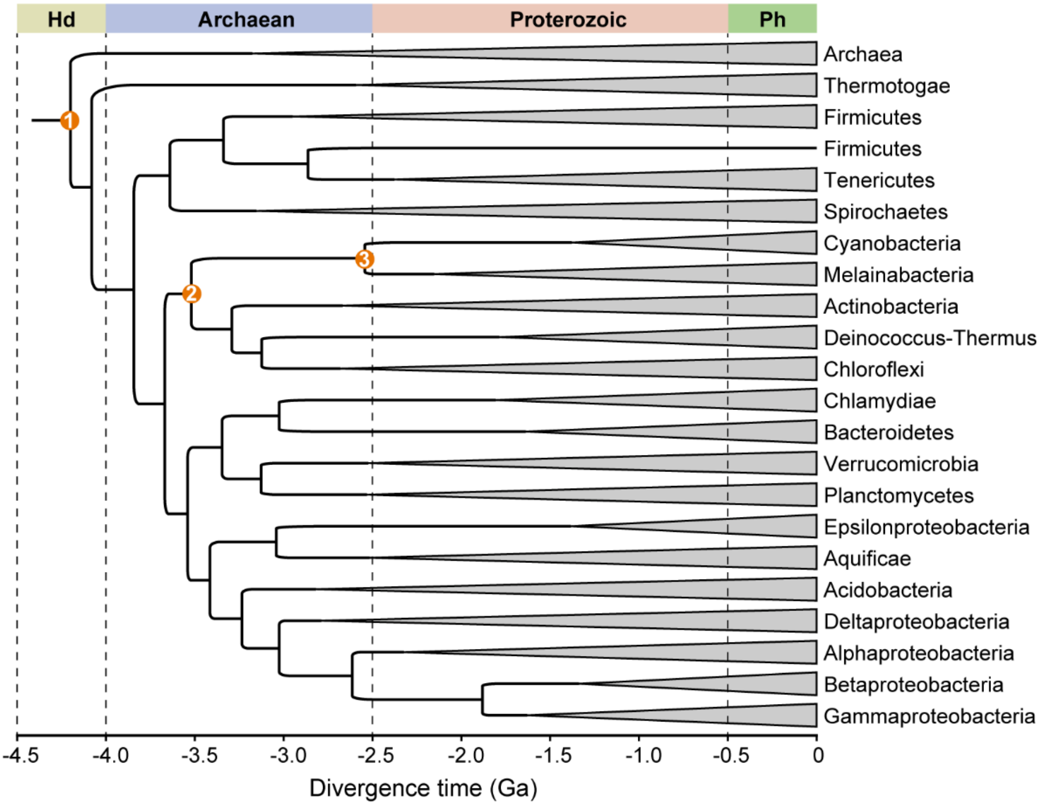


**Figure S17. Molecular dating of the phylogenetic tree of bacteria.** Orange circles on the tree indicate the temporal constrains (node 1: < 4.52 Ga; node 2: > 3.225 Ga; node 3: 2.5-2.6 Ga).

**Table S1. Function of KOs correlated to the genomic GC content.**

| KO | Protein | Function in previous studies |
|---|---|---|
| K01523 | HisE | Involve in the biosynthesis of histidine (*1*) |
| K07684 | NarL | Function as a nitrate/nitrite response regulator (*2*) |
| K01496 | HisI | Involve in the biosynthesis of histidine (*1*) |
| K05838 | YbbN | Function as a chaperone of β-clamp to cope with heat stress (*3*) |
| K00982 | GlnE | Regulate the glutamine synthetase in response to nitrogen limitation (*4*) |
| K00528 | FNR | Participate in the defense against oxidative damage (*5*) |
| K00697 | OtsA | Involve in the synthesis of trehalose in response to heat, cold and osmotic stresses (*6*) |
| K14998 | SURF1 | Regulate the assembly of cytochrome c oxidase involved in aerobic respiration (*7*) |
| K00970 | PcnB | Accelerate the mRNA degradation according to growth conditions (*8*) |
| K20712 | GlnA | Participate in the biodegradation and metabolism of xenobiotics (*9*) |
| K14267 | DapC | Involve in the biosynthesis of lysine (*10*) |
| K06980 | YgfZ | A tRNA-modifying protein which can increase the resistance to oxidative stress (*11*) |

| | | |
|---|---|---|
| K20035 | DmdC | Involve in the assimilation of dimethylsulphoniopropionate produced by marine phytoplankton (*12*) |
| K06117 | | Involve in the recycle and catabolism of glycerophospholipid (*9*) |
| K11755 | HisIE | Involve in the biosynthesis of histidine (*1*) |
| K21562 | FLP | Function as an anaerobic regulatory protein in response to oxidative stress (*13*) |
| K18682 | Rny | Interact with glycolytic proteins by processing the mRNA of *gapA* operon (*14*) |
| K15460 | YfiC | Modify valine-specific tRNA to promote growth in combating hyperosmotic and oxidative stress (*15*) |
| K00588 | | Involve in the biosynthesis of secondary metabolites (*16*) |
| K03495 | MnmG | Involve in the modification of tRNAs (*17*) |
| K01258 | PepT | Catalyze the release of free amino acids for nutritional utilization of tripeptides (*18*) |
| K03650 | MnmE | Involve in the modification of tRNAs (*17*) |
| K18672 | DacA | Convert ATP or ADP into the c-di-AMP which regulate various cellular processes (*19*) |
| K00243 | | Uncharacterized protein |
| K00974 | CCA | Add the nucleotides CCA onto the 3’ end of tRNA precursors (*20*) |
| K01493 | ComEB | Involve in the uptake of exogenous DNA under nutrient |

| | | |
|---|---|---|
| | | starvation (*21*) |
| K01893 | AsnRS | Catalyze the specific aminoacylation of tRNA Asn with asparagine (*22*) |
| K00850 | PfkA | Be required for the utilization of more carbon sources (*23*) |

**Data S1. (separate file)**

List of the bacterial representative genomes analyzed in the present study.

**Data S2. (separate file)**

List of the identified cold-adapted bacterial species analyzed in the present study.

**Data S3. (separate file)**

Chemical properties of 20 proteinogenic amino acids.
